# Comparison of SARS-CoV-2 variants of concern in primary human nasal cultures demonstrates Delta as most cytopathic and Omicron as fastest replicating

**DOI:** 10.1101/2023.08.24.553565

**Authors:** Nikhila S Tanneti, Anant K Patel, Li Hui Tan, Andrew D Marques, Ranawaka A P M Perera, Scott Sherrill-Mix, Brendan J Kelly, David M Renner, Ronald G Collman, Kyle Rodino, Carole Lee, Frederic D Bushman, Noam A Cohen, Susan R Weiss

## Abstract

The SARS-CoV-2 pandemic was marked with emerging viral variants, some of which were designated as variants of concern (VOCs) due to selection and rapid circulation in the human population. Here we elucidate functional features of each VOC linked to variations in replication rate. Patient-derived primary nasal cultures grown at air-liquid-interface (ALI) were used to model upper-respiratory infection and human lung epithelial cell lines used to model lower-respiratory infection. All VOCs replicated to higher titers than the ancestral virus, suggesting a selection for replication efficiency. In primary nasal cultures, Omicron replicated to the highest titers at early time points, followed by Delta, paralleling comparative studies of population sampling. All SARS-CoV-2 viruses entered the cell primarily via a transmembrane serine protease 2 (TMPRSS2)-dependent pathway, and Omicron was more likely to use an endosomal route of entry. All VOCs activated and overcame dsRNA-induced cellular responses including interferon (IFN) signaling, oligoadenylate ribonuclease L degradation and protein kinase R activation. Among the VOCs, Omicron infection induced expression of the most IFN and IFN stimulated genes. Infections in nasal cultures resulted in cellular damage, including a compromise of cell-barrier integrity and loss of nasal cilia and ciliary beating function, especially during Delta infection. Overall, Omicron was optimized for replication in the upper-respiratory system and least-favorable in the lower-respiratory cell line; and Delta was the most cytopathic for both upper and lower respiratory cells. Our findings highlight the functional differences among VOCs at the cellular level and imply distinct mechanisms of pathogenesis in infected individuals.

**Importance:** Comparative analysis of infections by SARS-CoV-2 ancestral virus and variants of concern including Alpha, Beta, Delta, and Omicron, indicated that variants were selected for efficiency in replication. In infections of patient-derived primary nasal cultures grown at air-liquid-interface to model upper-respiratory infection, Omicron reached highest titers at early time points, a finding that was confirmed by parallel population sampling studies. While all infections overcame dsRNA-mediated host responses, infections with Omicron induced the strongest interferon and interferon stimulated gene response. In both primary nasal cultures and lower-respiratory cell line infections by Delta were most damaging to the cells as indicated by syncytia formation, loss of cell barrier integrity and nasal ciliary function.

## INTRODUCTION

The SARS-CoV-2 pandemic has been marked by evolution of the ancestral virus strains, Wuhan in the east and Washington in the west, into new variants. The World Health Organization has identified some of the variants that pose an increased risk for global public health as variants of concern (VOCs). Characteristics for VOCs include an increase in virus transmissibility and virulence, and/or a decrease in response to current vaccines, public health measures and therapeutics (1).

Four major VOC have emerged over the course of the pandemic. The Alpha VOC, B.1.1.7 pango lineage virus, was first documented in the United Kingdom in September 2020. The Beta VOC, B.1.351 pango lineage virus, was first documented in South Africa in May 2020. The Delta VOC, B.1.617.2 pango lineage virus, was first identified in India in October 2020. The B.1.1.529 pango lineage Omicron variant was first documented in South Africa in November 2021 (2); however, more recent reports suggest an earlier emergence. Since then, several sub-variants have emerged from the Omicron lineage, including the BA.5 and XBB.1.5 which have quickly established global dominance, and finally BQ.1 in some regions (3).

Clinical retrospective studies report differences in patient outcomes among VOCs. Compared to the ancestral strain, patients infected with Delta and Alpha variants experienced heightened disease severity, such as increase in oxygen requirement, longer hospitalization and morbidity (4). Patients infected with the Omicron variant were less likely to develop severe COVID-19, require hospitalization, and had lower rates of in-hospital mortality compared to Delta-infected patients but the assessment of these traits was confounded by the concomitant development of immunity from prior exposure and vaccination in these populations (5, 6).

Variant-specific mutations to the ancestral SARS-CoV-2 genome are of interest for their potential roles in facilitating virus pathogenesis and spread. Many of the amino acid substitutions are found in the spike-encoding region of the SARS-CoV-2 genome. The spike (S) protein, composed of S1 and S2 subunits, binds to the host angiotensin converting enzyme 2 (ACE2) receptor to facilitate virus-host membrane fusion, virus entry, and modulates host immune responses (7-9). Several studies attribute the enhanced immune evasion by the variants to substitutions in the spike protein. Amino acid substitutions that render the furin protease recognition site at the spike S1/S2 subunit junction more basic have been associated with enhanced viral fitness for the Delta VOC, and to a lesser extent for the Alpha and Omicron variants (10). Deletions and substitutions within and adjacent to the furin protease recognition site have been associated with virus attenuation (11, 12). The role of mutations outside of the spike gene, although not as well characterized, likely also contribute to differences in pathogenesis. For example, mutations in the nucleocapsid gene have been associated with increased virus replication and pathogenesis (13).

To assess genome-wide differences between SARS-CoV-2 VOCs in a controlled system, we compared molecular replication mechanism among full length, replication competent Alpha, Beta, Delta and Omicron VOCs to the ancestral Washington (WA1). All viruses were compared for replication kinetics and cellular responses to infections in patient-derived primary nasal cultures, with the goal of modelling the first step of infection. In addition, all experiments were repeated in cell lines derived from human lung tissues, including Calu-3 and A549, to facilitate more mechanistic studies that are not limited by primary cells. The cell lines also provide the context of a lower-respiratory model for infections. Our experiments clarify differences among variants in virus entry, virus replication, cell-to-cell spread of the virus, and activation of host innate immune responses and antiviral pathways.

This study adds to our understanding of SARS-CoV-2 variants in several ways. We compared the ancestral WA1 to Alpha, Beta, Delta and Omicron variants while many focus on a subset of this group. All experiments involve infections with authentic viruses in a BSL3 facility rather than the use of pseudoviruses or protein expression systems. In addition to quantifying RNA copy numbers, which can be misleading with RNA viruses, our experiments also quantify infectious viruses. While several useful animal models have been developed to study SARS-CoV-2 (14, 15) we find that our patient-derived nasal epithelia model more faithfully reflects replication kinetics in the human nose, the first site of infection.

## RESULTS

### VOCs are selected for increased replication in upper respiratory cells

SARS-CoV-2 infections are initiated in the upper respiratory system, so we sought to model this step by comparing infections in primary cells collected from nasal cavities of patients undergoing rhinologic evaluation. These cells were cultured on transwells at an air-liquid-interface (ALI) to recapitulate the natural state of the nasal epithelium, as we have reported previously (16). In virus growth curve assays, all SARS-CoV-2 viruses replicated to high titers (> 1x10^5 PFUs/mL), and all VOCs replicated to higher titers than WA1 (**Fig 1A**). Omicron reached the highest titers at early time points of infection, starting at 24 hours post infection (hpi), and maintained the highest titer until 72 hpi, after which titers start to drop. WA1 generally produced the lowest titers compared to all variants. These results suggest that emerging variants have been selected for replication in human nasal cultures, as Delta and Omicron, the two most recent VOCs, reached the highest titer. The growth curve experiment was ended at 96 hpi as the infected nasal cultures began to die and titers were no longer significantly different at this point.

**Figure 1:**
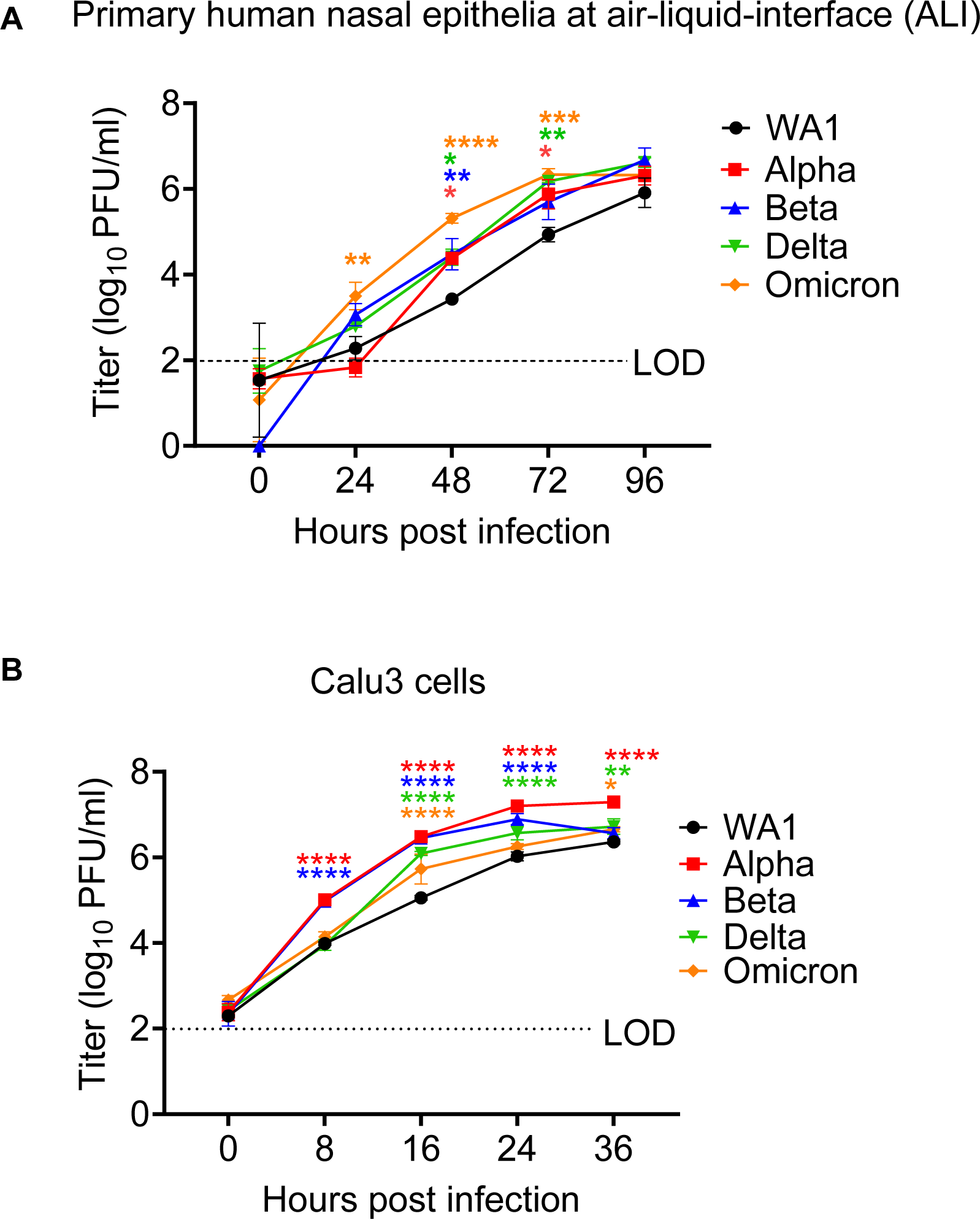
Replication of SARS-CoV-2 WA1 and VOCs. (A&B) Human primary nasal epithelia ALI cultures (A) or Calu-3 cells (B) were infected with SARS-CoV-2 WA1, Alpha, Beta, Delta and Omicron at MOI 0.1 PFU/cell, and apically shed virus was titered by plaque assay at indicated hours post infection. (A) Titers from nasal cultures are an average of 3 independent experiments, each experiment was performed with three individual donor cells totaling to 9 individual donors per virus. (B) Titers from Calu3 are an average of two independent experiments each in triplicates. (A and B) Graphed values represent mean with standard deviation, and statistics for both A and B were performed with ordinary two-way ANOVA comparing VOCs versus WA1 within a time point, multiple comparisons adjusted P-values: *P<0.01, **P<0.001, ***P<0.0001, ***P<0.00001.

Coronavirus infections naturally progress from the upper to the lower respiratory system. To investigate the relative efficiencies of replication of the variants in cells of the lower respiratory system, we used the Calu3 cell line derived from human lung epithelia. Similar to observations in the primary nasal cultures, all VOCs replicated to higher titers than the ancestral WA1 in Calu3, suggesting that all VOCs were selected for more efficient replication than the ancestral SARS-CoV-2 (**Fig 1B**). However, the Alpha and Beta VOCs reached significantly higher titers than Omicron (at 8 hpi) in Calu3 infections while Omicron reached significantly higher titers than Alpha/Beta/Delta in nasal cultures. Also, all viruses reached peak titers at an earlier time (24-36 hpi) in Calu3 cells than in nasal cultures. As the Calu3 cells begin to show signs of cytopathic effect and cell death around 36 hpi, the experiment could not be extended to further duration. Together these results suggest that while all VOCs replicate more efficiently than the ancestral virus, Omicron is especially selected for heightened replication in nasal cultures.

To ensure the integrity of the experiments, genomic RNA from each virus stock was sequenced and aligned against the Washington A reference genome before further analysis (**Fig S1**). The alignment confirmed that all viruses used in this study maintain the defining mutations of each lineage. Additionally, it confirmed that no new substitutions have become fixed at the known hotspots, including the furin protease recognition site (11).

The COVID-19 pandemic exhibited waves of illnesses in the colder months, suggesting a seasonal pattern similar to other respiratory viruses (17). Replication fitness at different temperatures may impact the potential for VOCs to cause seasonal outbreaks. To investigate whether the VOCs have adapted to colder temperatures, similar to the seasonal common cold coronaviruses, infections with WA1, Delta and Omicron viruses were performed in nasal cultures at 33°C and 37°C (**Fig S3A**). We did not observe any significant differences in titer when comparing infections at both temperatures up to 96 hpi, suggesting that SARS-CoV-2 replication is not temperature sensitive early in infection. However, we (16) and others have reported preference of WA1 at 33°C at later times post infection.

### The influence of furin protease cleavage of spike on viral entry and cell-to-cell spread

To understand the mechanisms of increased replication of the VOCs, cell to cell spread of the virus was investigated. Cleavage of the spike protein at the furin recognition site facilitates both virus entry and cell-to-cell virus spread, another mechanism that could contribute to increased replication of the variants. The basic amino acids at the furin recognition site of the spike protein, PRRAR in WA1, recruit proteases necessary for the cleavage at the S1/S2 junction (18). This region is sensitive to substitutions which influence infection and replication (11, 19-21). We hypothesized that the amino acid substitutions in VOCs that render the furin cleavage site more basic would be more efficiently cleaved, leading to increased cell-to-cell spread. Specifically, we hypothesized that Delta, which encodes the most basic amino acids in the cleavage-site (RRRAR), will generate the most cleaved spike, followed by Alpha and Omicron (both HRRAR) generating equal levels of cleaved spike. Beta and WA1, which encode S1/S2 site with the least basic charge (PRRAR), were expected to generate the least cleaved spike.

To measure the ratios of cleaved versus full-length spike protein levels among VOCs, a western blot was performed using protein lysates from infected primary nasal cultures to assess the ratios of cleaved spike (**Fig 2A**). As predicted, quantification of the cleaved spike from three separate donors showed that Delta infections generated the highest proportion of cleaved spike (71%), followed by Alpha infection (52%), Omicron and Beta infections generated similar levels of cleaved spike (∼44% each), and the lowest by WA1 infection (32%) (**Fig 2B**). The elevated levels of cleaved spike in infections by Alpha and Delta compared to WA1 are consistent with the idea that these VOCs have evolved to optimize spread in human nasal cells. However, infection with Omicron produced the highest titers **(Fig 1A)** despite inefficient spike cleavage (**Fig 2A-B**), indicating the importance of factors affecting other replication steps.

**Figure 2:**
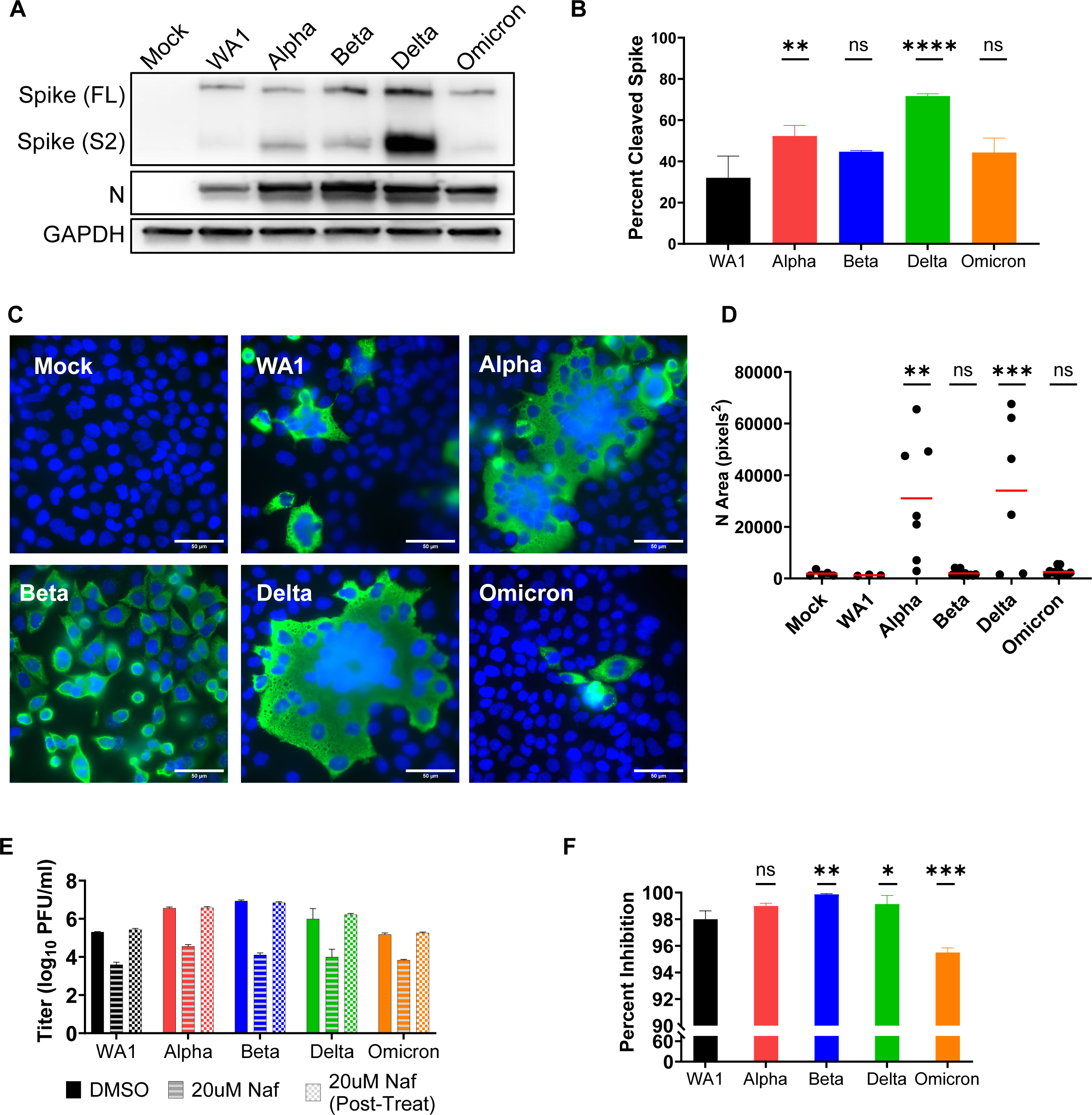
Virus entry and spread. (A) Protein lysates from nasal cultures infected with SARS-CoV-2 WA1 and VOCs were analyzed by polyacrylamide gel electrophoresis followed by immunoblotting with antibodies against SARS-CoV-2 proteins, nucleocapsid (N) and spike S2, which recognizes the full length (FL) and cleaved (S2) forms. Cellular protein GAPDH was used for loading control of the gels. Western blot depicted is representative of 3 independent donor infection. (B) Percent cleaved spike was calculated using the following formula: S2 / (FL + S2). Graphed values are an averaged from 3 independent western blots and statistics were performed with ordinary one-way ANOVA comparing VOC against WA1, adjusted P-values: **p<0.001, ****p<0.00001. (C) A549^ACE2^ cells infected with SARS-CoV-2 WA1 and VOCs (MOI=0.01 for 24hpi), fixed and stained with fluorescently labeled antibodies DAPI (blue) and SARS-CoV-2 nucleocapsid (green). Images are representative of 3 independent infections. (D) Independent clusters expressing N were quantified for area (pixel^2^) from 3 independent experiments. Graph represents individual values with mean in red, statistics were performed with two-way ANOVA with multiple comparisons of VOCs versus WA1, adjusted P-values: **p<0.001, ***p<0.0001. (E) Calu3 cells were mock treated (DMSO), pre-infection treated with 20uM Nafamostat (Naf) for 2 hours, or pre- and post-infection treated with 20uM Naf for 2 hours. Infections with SARS-CoV-2 WA1 and VOCs were performed at MOI=0.1 and shed virus was collected at 16 hpi for titer by plaque assay. Graphed values are an average of 2 independent infections each performed with biological triplicates. (F) Percent inhibition in virus titer after Naf treatments. Graphed values represent mean with standard deviation, and statistics were performed with ordinary one-way ANOVA comparing the VOC against WA1, adjusted P-values: *p<0.01, **p<0.001, ***p<0.0001.

Similar experiment in the VeroE6^TMPRSS2^ cell line yielded different relative ratios of cleaved spike among VOCs (**Fig S2A**). In this western blot, protein lysates from infected VeroE6^TMPRSS2^ cells were analyzed, using equal levels of full-length spike to visualize differences in the ratios of cleaved spike among the VOCs. While the highest ratio of cleaved spike was produced with Delta infection (67% cleaved) as predicted and observed by others (22), contrary to our hypothesis, infection with the Beta generated the second most cleaved spike. Infections with Alpha and Omicron VOC generated different levels of cleaved spike despite sharing the same furin cleavage site sequence, similar to observations of spike proteins from primary nasal cultures. These results add to our understanding that while the strong basic charge at the furin cleavage facilitates spike cleavage, there are other factors beyond the PRRAR sequence that facilitate cleavage. In addition, cell type-specific biology, e.g. variation in the abundances of various proteases, may influences the cleavage of spike. As validation of our spike cleavage assay we included infections with a positive control virus (icWT), expressing a spike containing the ancestral PRRAR sequence and a negative control virus (icΔPRRA) expressing a spike with a deletion of the PRRA sequence, both generated with the infectious clone reverse genetics system (**Fig S2A**) (11). As expected, infections with both WA1 and icWT, both encode a PRRAR furin recognitions site and generate comparable ratios of S2 spike, while infection with the icΔPRRA expressing a spike lacking the PRRA sequence did not generate any cleaved spike (**Fig S2A**).

An immunofluorescence staining assay (IFA) was used to assay spread of infection. A monolayer of A549^ACE2^ cells were infected at a low MOI and stained with nucleocapsid and DAPI to visualize syncytia. Compared to WA1, Alpha and Delta infections generated strikingly larger syncytia with the classical clustering of nuclei in the middle (**Fig 2C**); syncytia size was quantified by measuring the area of nucleocapsid positive (green) clusters **(Fig 2D)**. The formation of syncytia, especially during Delta infection was also observed in primary human nasal cultures (**Figs 7C-D** and **S3B-C**); however, the heterogenous nature of primary nasal cultures and the weakly adhering nature of Calu3 cells do not make them ideal for visualizing syncytia. These data suggest that the Alpha and Delta VOCs are efficient at cell-to-cell spread, likely due to efficient spike cleavage and high fusogenic activity. However, cell-to-cell spread of virus by syncytia formation does not correlate with rate of replication, suggesting the role of different pathways driving these mechanisms.

### SARS-CoV-2 viruses enter primarily by the transmembrane protease serine 2 (TMPRSS2)-mediated plasma membrane fusion pathways

During virus entry the SARS-CoV-2 spike protein is cleaved by TMPRSS2, a serine protease which activates the spike to initiate fusion at the plasma membrane (23-25). It is thought that viruses that express spike with an additional basic amino acid(s) in the furin cleavage recognition site than WA1 (Alpha, Delta, Omicron) would, be likely to use the TMPRSS2 dependent plasma membrane route of infection. However, there are reports of Omicron entering cells in a TMPRSS2-independent endosomal route in some cell types(26-30). To investigate entry via the plasma membrane or endosomal routes, we used the protease inhibitor Nafamostat that blocks TMPRSS2 and inhibits entry by this pathway. Upon treatment with Nafamostat we observed a large reduction in virus titer for all viruses (**Fig 2E**). Compared to untreated controls (DMSO) virus titer was inhibited by 95-99% for all viruses (**Fig 2F**), suggesting that all VOCs primarily use the plasma membrane as the major route of entry. Omicron titer was affected the least, suggesting that Omicron is less dependent on the TMPRSS2-mediated entry pathway compared to the other SARS-CoV-2 viruses, and therefore may use the alternative endosomal route more than other VOCs. Similar results have been reported by others (24, 31-33). However, it is important to note that while Omicron is the least dependent VOC on the plasma membrane fusion pathway, it still enters primarily by this pathway in Calu3 cells.

### SARS-CoV-2 WA1 and VOC plaque morphology

Quantification of infectious virus titer was performed in VeroE6 cells by a plaque assay, during which we observed a variation in plaque morphology and size (**Fig 3A-B**). WA1 generated the largest plaques (average 1542 px^2^) but also displayed the most range in plaque area, as reported by others (34). While all VOCs generated smaller plaques than WA1, the plaques generated with Alpha and Delta VOCs were strikingly smaller (average of 211 px^2^ and 231 px^2^, respectively). However, it is worth noting that differences in plaque area are abrogated on VeroE6^Tmprss2^ overexpressing cell lines, as reported by others, suggesting a role of cellular protease levels on plaque morphology (35, 36). It is intriguing that, in comparison to WA1, Alpha and Delta generate smaller plaques even though they produce higher titers of virus in growth curves (**Fig 1A-B**) and form larger syncytia (**Fig 2C-D**). This demonstrates that for SARS-CoV-2, plaque size does not necessarily correlate with replication and cell-to-cell spread, an observation made previously about murine coronavirus strains (37).

**Figure 3:**
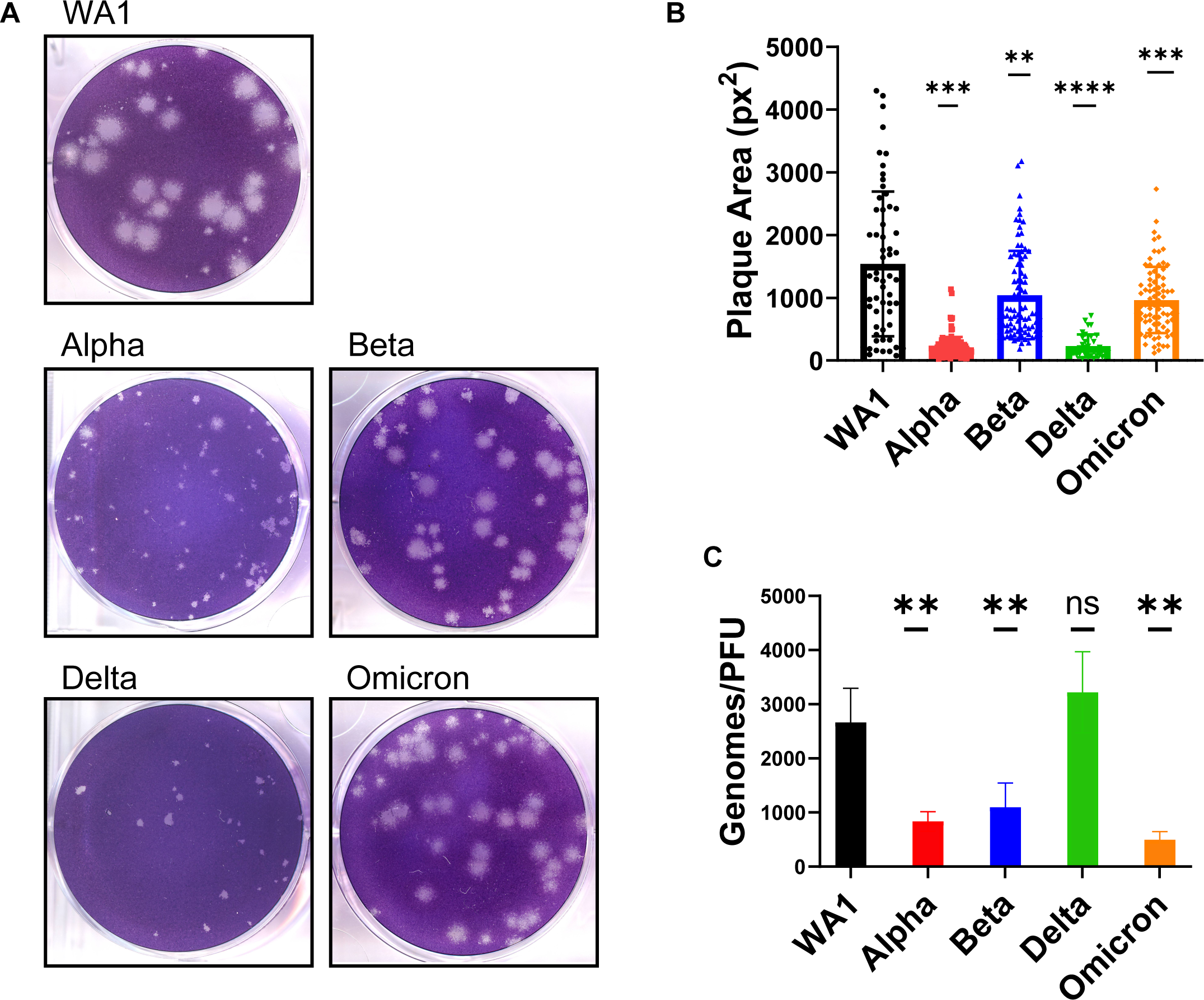
Comparison of plaque size and genome/PFU ratio among WA1 and VOCs. (A) Plaque assay of SARS-CoV-2 viruses was performed on VeroE6 cells. (B) Three independent plaque assays per virus were quantified for plaque area (pixel^2^), and graph represents the mean of individual plaque areas with standard deviation. Statistics were performed with ordinary two-way ANOVA comparing VOCs versus WA1, adjusted P-values: **P<0.001, ***P<0.0001, ****P<0.00001. (C) The virus in supernatant collected from Calu3 cells infected at MOI=0.1 at 24 hpi was assessed for genomes using qRT-PCR with primers specific for SARS-CoV-2 RdRp and compared to PFUs quantified by plaque assay. Values are an average of 3 independent experiments, each performed in triplicate. Graph represents mean with standard deviation, and statistics were performed with ordinary one-way ANOVA comparing VOCs versus WA1, adjusted P-values: **P<0.001.

The differences in infectious virus production in titer assays, despite comparable levels of intracellular genome replication by RT-qPCR (**Fig S2B**), prompted us to quantify the particle-to-PFU ratio among VOCs. Compared to the WA1 virus, all VOCs, except Delta, secreted significantly fewer genomes per infectious virus (**Fig 3C**), suggesting that Alpha, Beta and Omicron VOCs are more efficient at producing infectious virus. The Delta VOC genome-to-PFU ratio was not significantly different from WA1.

### Omicron infection activates highest levels of interferon and interferon stimulated genes

dsRNAs generated during the replication of RNA viruses, including coronaviruses, are detected by host cells and elicit antiviral responses. There are three major cytosolic sensors of dsRNAs that induce innate immune responses (38). Detection of coronavirus dsRNA by MDA5 leads to induction of type I and type III interferons (IFN) and activation of interferon stimulated genes (ISGs), many of which encode proteins with antiviral activities. Sensing of dsRNA by oligoadenylate synthetases (OASs) leads to production of 2’,5’-oligoadenylates, which activate host ribonuclease (RNase) L to degrade host and viral single-stranded (ss)RNA. Activation by protein kinase R (PKR) leads to dimerization and autophosphorylation followed by phosphorylation of p-eIF2α and inhibition of translation. Both RNase L and PKR activation lead to reduced virus replication, apoptosis and inflammation (39). Thus, differential responses to these pathways are a potential explanation for the differences in pathogenesis of the VOCs.

To compare the induction of the interferon (INF) pathway among the SARS-CoV-2 viruses, reverse transcriptase-quantitative polymerase chain reaction (RT-qPCR) was used to quantify gene expression of type I IFN (*IFNB*), type III IFN (*IFNL1*), and ISGs, including 2’-5’-oligoadenylate synthetase 2 (*OAS2)* and interferon induced protein with tetratricopeptide repeats 1 (*IFIT1)* in RNA from infected cells (**Fig 4**). Host responses to infection were measured in both primary nasal cultures (**Fig 4A**) and Calu3 cells (**Fig 4B**) to understand if all viruses activate same host response in both cell types, or if there is a cell type or VOC dependent response. To ensure productive virus replication, genome copy levels were quantified from infected cell lysates, by performing RT-qPCR using primers to amplify nsp12 RNA dependent RNA polymerase, RdRp sequences. High genome copies of RdRp confirmed that all viruses reached high and productive replication in both cell types. The overall ratios of gene expressions do not correlate with virus genome copy levels suggesting a reliable quantification of innate-immune responses induced in primary nasal cultures and Calu3 cells that is not skewed by viral RNA copy numbers.

**Figure 4:**
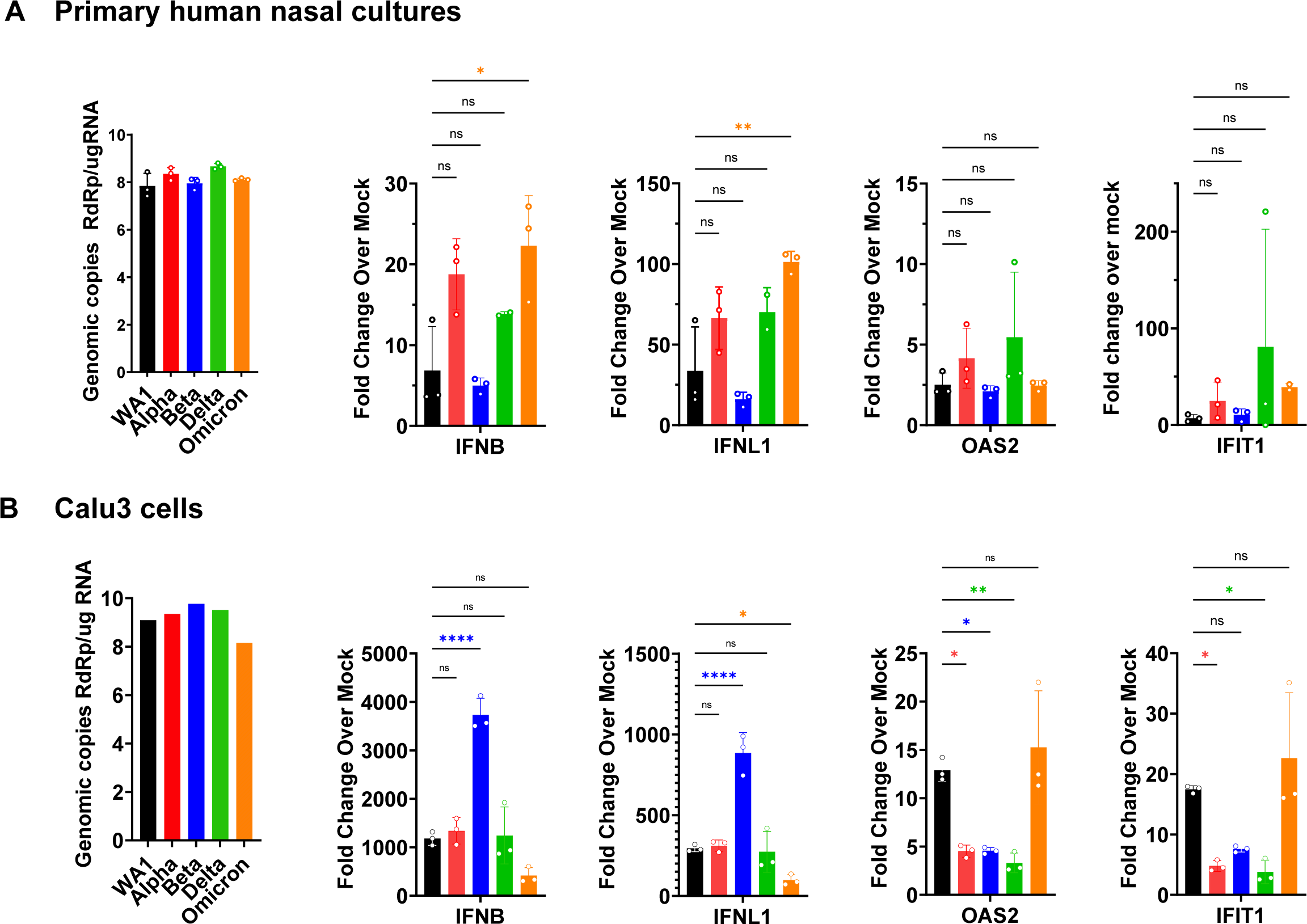
RT-qPCR of interferon and interferon-stimulated responses to SARS-CoV-2 infections. (A&B) Infections (MOI=0.1) with indicated viruses were performed in (A) primary nasal cultures infected for 96hpi and (B) Calu3 cells at for 32hpi. Cells were lysed for RNA extraction and viral genomes were quantified by RT-qPCR with primers specific for SARS-CoV-2 RdRp, and human *IFNB*, *IFNL1*, *OAS2*, and *IFIT1*. Black bars indicate WA1, red for Alpha, blue for Beta, green for Delta and orange for Omicron. For each cell type, graphs represent one of two independent experiments, each performed in biological triplicates. For primary nasal cultures each experiment included a pool culture of 3 donors. Graphed values are mean with standard deviation, and statistics were performed with ordinary one-way ANOVA comparing VOCs versus WA1, adjusted P-values: *P<0.01, **P<0.001, ***P<0.0001, ***P<0.00001.

We observed that while all viruses activated INFB and INFL1 expression above mock levels in primary nasal cultures, Omicron significantly induced these pathways (**Fig 4A**). Consistent with these data, previous studies comparing IFN induction between Delta and Omicron infections in cell lines have also reported greater induction with Omicron (40, 41). However, we did not observe any differences by RT-qPCR for the ISGs, OAS2 and IFIT1, among the viruses (**Fig 4A**). In contrast, in Calu3 cells, Beta generated significantly greater IFN responses than the other viruses. However, induction of ISGs was still the highest with Omicron (**Fig 4B**). The discrepancies in gene induction between nasal cultures and Calu3 cells suggests cell-type dependent host responses, and in both contexts the infection was able to overcome cellular responses and lead to a productive replication.

To understand the activation of INF and ISGs on a broader scale we sequenced RNA from lysates of primary nasal cultures infected with each of the viruses (**Fig 5**). RNAseq data generated from infected and mock infected cells were analyzed for differentially expressed genes (DEGs) that exceed the threshold of log_2_ fold change greater than 1 and p-adjusted value less than 0.01 for hits with high significance. Infection with WA1 induced lowest number of DEGs while infection with Omicron induced the highest number of DEGs (16 and 823, respectively). Infection with Alpha and Beta also resulted in a relatively high number of DEGs (475 and 202, respectively) while Beta was relatively lower (23 DEGs). All DEGs were processed for ISG gene ontology, which yielded a striking number of ISGs associated with Omicron infection (51 ISGs), followed by Alpha infection (28 ISGs) (**Fig 5E and B**). While infections with Delta (23 ISGs), Beta (18 ISGs) and WA1 (14 ISGs) also generated an ISGs response, the number and amplitude of the upregulation was much lower than those induced by Omicron and Alpha infections. Overall, our data shows that Omicron infection induces the strongest IFN and ISGs responses.

**Figure 5:**
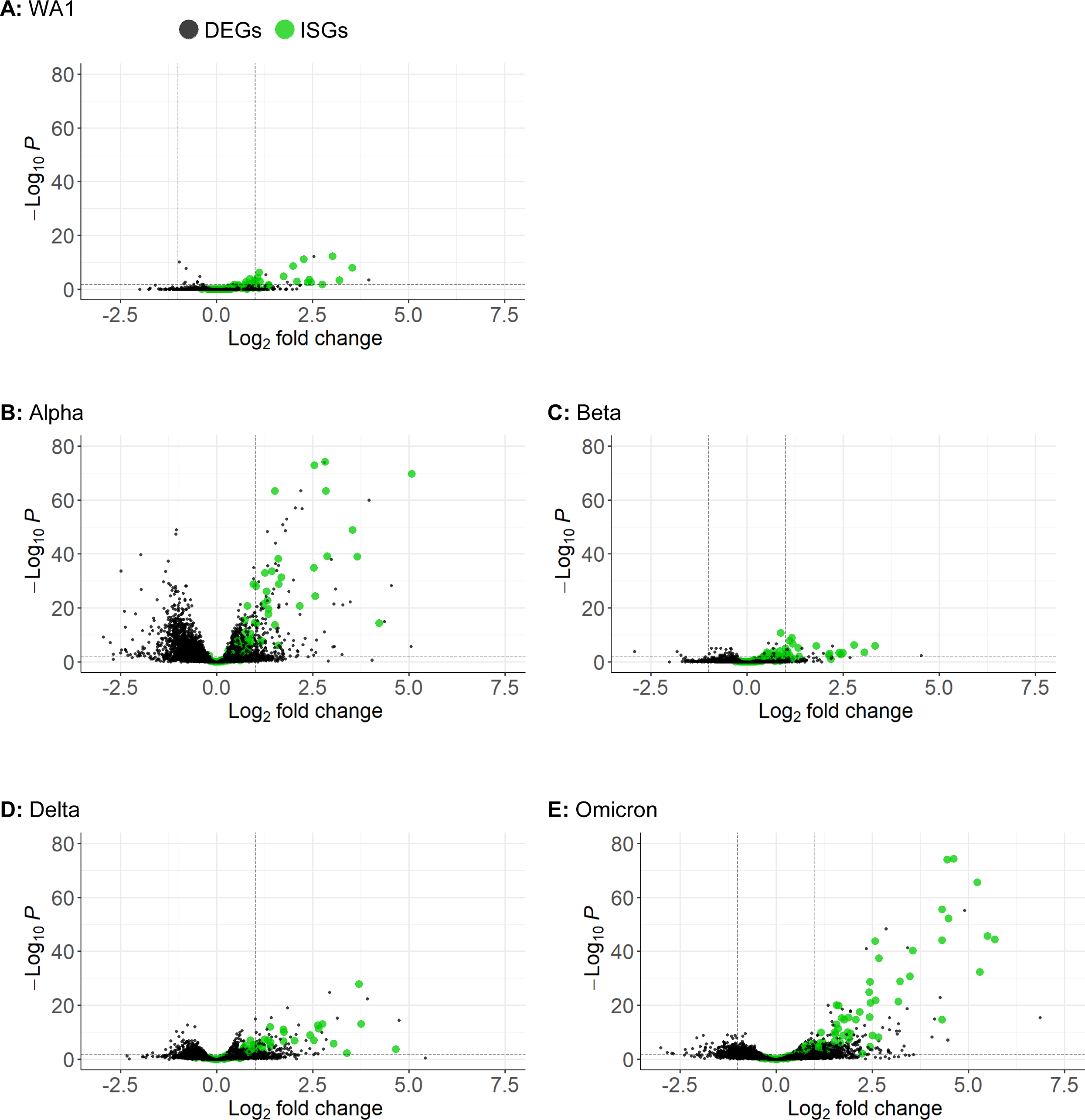
RNA-Seq of primary nasal cultures infected with SARS-CoV-2 WA1 and VOCs. (A-E) Primary nasal cultures pooled from 4 donors were infected with WA1 or VOCs at MOI=0.1 and RNA was extracted at 96 hpi for RNA-Seq analysis. Each volcano plot represents significantly differentially expressing genes (DEGs) for the indicated infection compared to mock-infected primary nasal cultures. Black dots indicate DEGs, and green dots highlight interferon stimulated genes (ISGs). The number of graphed variables per condition are (A) 13614, (B) 13385, (C) 13623, (D) 13452 and (E) and 13515.

### Double-stranded (ds)RNA-induced innate immune responses to SARS-CoV-2 variants

The activation of the PKR pathway during SARS-CoV-2 infection of primary nasal cultures and A549^ACE2^ cells was assessed by western blot (**Fig 6A-B**). Phosphorylated-PKR (p-PKR) was detected in lysates of nasal cultures infected with all strains (Fig 6A) and in in A549 ^ACE2^ cells for WA1, Alpha and Omicron (**Fig 6B**), above the level of mock-infected cells, indicating that SARS-CoV-2 WA1 and VOCs activate the PKR pathway. Phosphorylated-eIF2α was also detected during infection with all strains in A549 ^ACE2^ cells. However, the level of p-eIF2α over mock infected cells, was variable in nasal cultures, a reproducible observation for the nasal culture model, which may be due to the low percentage of infected cells and that eIF2α level is not IFN induced while PKR level as well as phosphorylation of PKR is IFN dependent. Together these results suggest that all SARS-CoV-2 infections in primary nasal cultures and A549^ACE2^ cells induce the PKR and eIF2α pathway (**Fig 6A-B**).

**Figure 6:**
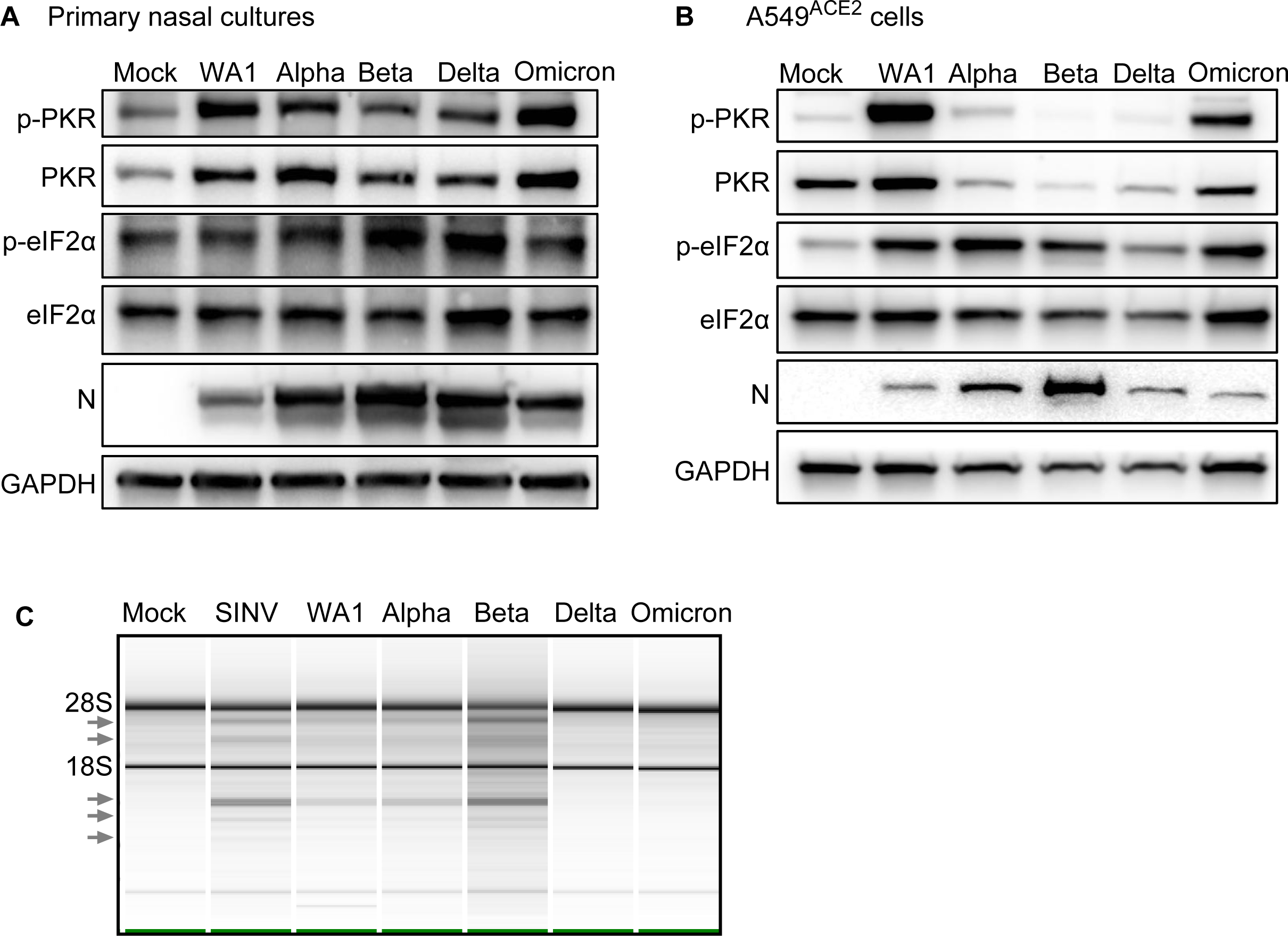
dsRNA-induced pathways during SARS-CoV-2 WA1 and VOCs infections. (A) Primary nasal cultures were infected at MOI=0.1. Cells were lysed at 96hpi, protein extracted and analyzed by western blot for phosphorylated PRK (p-PKR), PKR, phosphorylated eIF2α (p-eIF2α), eIF2α, SARS-CoV-2 nucleocapsid (N) and GAPDH. (B) A549^ACE2^ cells were infected at MOI=0.1 for 72hpi. Cells were lysed, protein extracted and analyzed by western blot for p-PKR, PKR, p-eIF2α, eIF2α, N and GAPDH. (A &B) Images are representative of western blots from 3 independent infections. (C) Infections were performed in A549^ACE2^ cells at MOI=0.1. At 48hpi (SARS-CoV-2 strains), and 24hpi (SINV), cells were lysed, RNA extracted and analyzed on a Bioanalyzer. Arrows indicate bands of degraded RNAs. Image is representative of two independent experiments.

The activation of the RNase L pathway can be inferred by assessing rRNA degradation using a Bioanalyzer (**Fig 6C**). Due to consistently undetectable levels of degradation of rRNA bands in lysates from primary nasal epithelial cells, possibly due to a low percentage of infected cells as above for p-eIF2α levels, we assessed lysates from infected A549^ACE2^ cells for rRNA degradation. Compared to mock-infected cell, the positive control SINV infection was associated accumulation of rRNA degradation products (arrows), as expected. Among the SARS-CoV-2 viruses, WA1, Alpha, and Beta generated rRNA degradation products. The reduced intensities of rRNA degradation in Delta and Omicron infection could be due to delayed progression of infection, as indicated by lower nucleocapsid (N) levels (**Fig 6C**) and titer (**Fig S2C**) in A549^ACE2^ cells, rather than a muted response by the RNase L pathway.

### Damage to upper-respiratory cells associated with Delta infection

Infection by SARS-CoV-2 can last for prolonged periods, raising the question of possible damage to infected tissue. To compare damage caused by the different variants we measured the transepithelial electrical resistance (TEER), which measures electrical resistance across the cell membranes that is maintained by intact cell-to-cell junctions. As the nasal cultures become more confluent and differentiate, the TEER values rise; however, any compromise to cell-to-cell barrier integrity leads to loss of ion transport across the membrane which can be measured as a reduction of TEER value. We observed that while all VOCs display an initial increase in TEER (0-48 hpi) there is a significant drop in TEER during late infection (48-96hpi) (**Fig 7A**). The most dramatic and significant loss of TEER was observed during Delta infection, suggesting that Delta induces significantly more damage to cell-to-cell barrier integrity than other variants.

**Figure 7:**
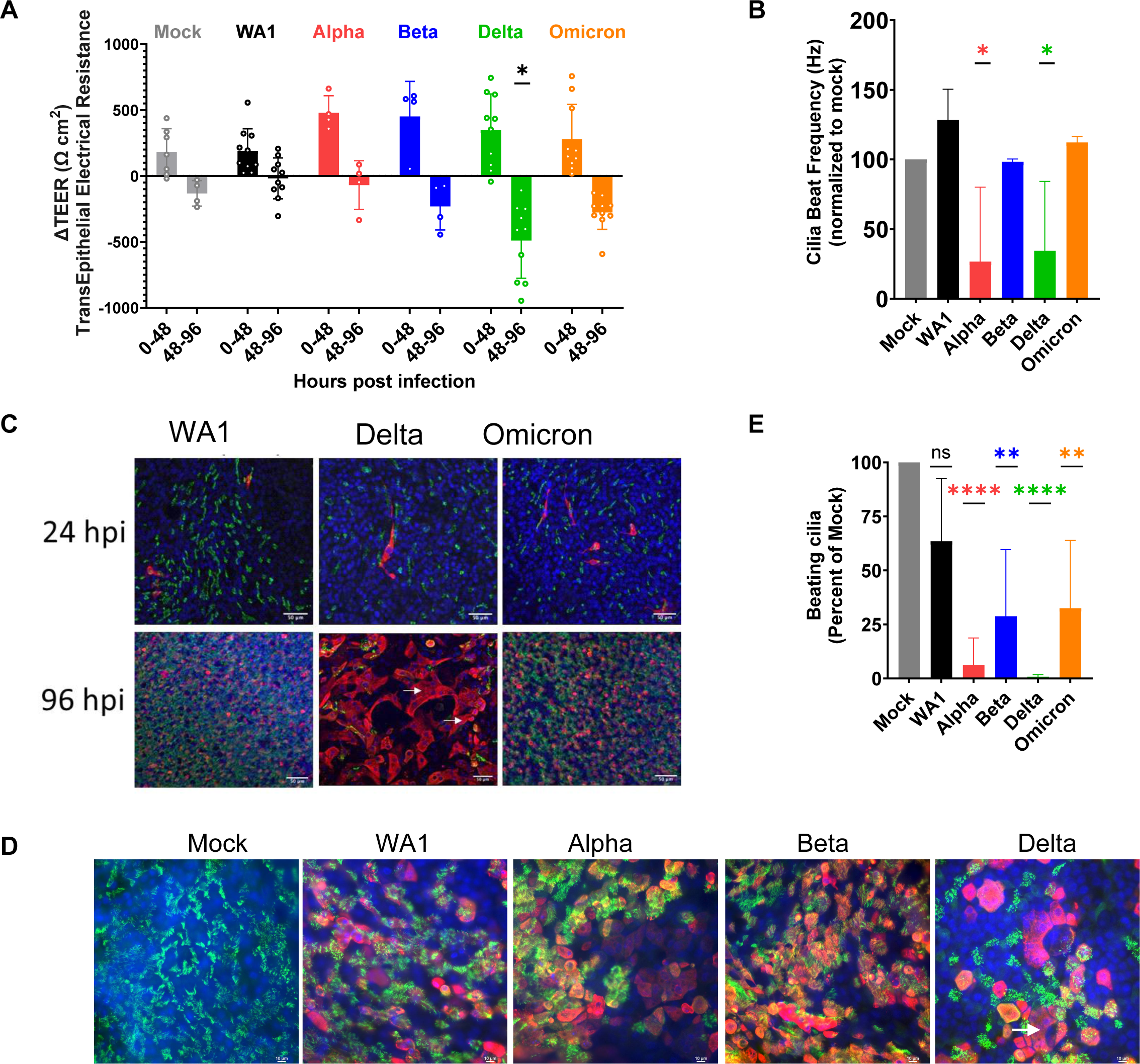
Infection with SARS-CoV-2 VOCs damages primary nasal cultures. Primary nasal epithelial cultures were infected with SARS-CoV-2 viruses at MOI=0.1 and processed at 96hpi for various assays. (A) Transepithelial electrical resistance (TEER) was quantified during SARS-CoV-2 infection of nasal cultures at 0, 48 and 96hpi. Values represent the difference in TEER (ΔTEER) during the indicated time span, and averaged from 3 independent experiments. Graphed values are mean with standard deviation, and statistics were performed with ordinary two-way ANOVA comparing VOCs versus WA1 within a time span, multiple comparison adjusted P-value: *P<0.01. (B) Ciliary beat frequency was measured in mock and infected nasal cultures at 96hpi. (C and D) Mock and infected nasal cultures from separate donors were fixed at 96hpi for immunofluorescence and confocal imaging. Antibodies were used to label DAPI (blue), β-tubulin (green) and SARS-CoV-2 nucleocapsid (red). Arrows indicate syncytia-like clusters. Scale bars indicate 50um (C) and 10um (D). Images are representative of infections in multiple donors. (E) Real-time videos of mock and infected nasal cultures at 96hpi were quantified for the area of actively beating cilia. (B and E) Graphed values represent mean with standard deviation and statistics were performed with one-way ANOVA using Dunnett’s multiple comparisons test for all infected conditions against mock/uninfected, with adjusted P-values: *P<0.05, **P<0.005, ****P<0.00005.

The nasal cultures include epithelial cells with projections of cilia on the apical surface, which facilitate transport and clearance of mucus that is generated in the nasal cavity. Beat frequency can be measured to assay activity and function of cilia as well as overall health of nasal cultures. Compared to uninfected mock cells, nasal cultures infected with Alpha and Delta displayed a notable 75% loss in ciliary beating frequency (**Fig 7B**). This suggests that SARS-CoV-2 infection compromises ciliary beating function in the nasal cell cultures, with different potency among variants.

Confocal microscopy was used to visualize the spread of infection and possible cell to cell fusion. During early infection (24 hpi) we observed that infected cells colocalized with the ciliary marker β-tubulin confirming that SARS-CoV-2 infects ciliated cells (**Fig 7C**) (16). Additionally, only a fraction (<5%) of the ciliated cells were infected at this early time point. During late infection (96 hpi) we observed a vast spread of infection throughout the nasal cultures. Notably, Delta infected nasal cultures showed some small syncytia-like clusters (**Fig 7C**, arrow), although not as pronounced as observed in lower respiratory cell lines (**Fig 2C**). The loss of the cytoskeletal marker, phalloidin, between cells in this cluster is further evidence of the formation of syncytia (**Fig S3C**). At higher magnification we observed that during later infection (96 hpi), Delta infected cells were negative for staining for β-tubulin, suggesting deciliation of the infected cells (**Fig 7D**). Using live microscopy and point-analysis, we also observed a reduction in the area where beating cilia could be detected (**Fig 7E**), further documenting deciliation. Together these results suggest that among the SARS-CoV-2 viruses, Delta VOC was the most cytopathic among WA1 and the other VOCs in human nasal epithelia.

### Population surveillance for SARS-CoV-2 reveals Delta and Omicron as fastest replicating

Given our goal of understanding molecular correlates of differences among variants, and the results described above, we investigated whether infection with different SARS-CoV-2 variants were associated with higher viral RNA levels as reported by RT-qPCR assays on clinical specimens. Samples were obtained from a program that paired viral whole genome sequencing with collection of clinical metadata, so that relative viral RNA levels were linked to the variant calls (42). Samples were analyzed from 2,722 infected participants from the Delaware Valley region. Our analysis pools Ct values from different analytical technologies that were in place based on the location of sample collection and the stage of the epidemic (**Fig S1B**). Due to the low prevalence of Beta infection in the Delaware Valley region our data does not include this VOC. Data from several sampling and analysis pipelines were combined, so a Bayesian analytical framework was used to control confounders and assess the significance of differences among variants (**Fig 8**). Relative abundance was quantified as cycle of threshold (Ct) in the qPCR, so that lower values indicate higher abundances. It is worth noting that some variants were associated with overwhelmed healthcare systems which might have resulted in longer wait times for testing, which could influence Ct values for some data points. Additionally, data for testing time since infection was not available, so Ct cannot be adjusted for days since infection.

**Figure 8:**
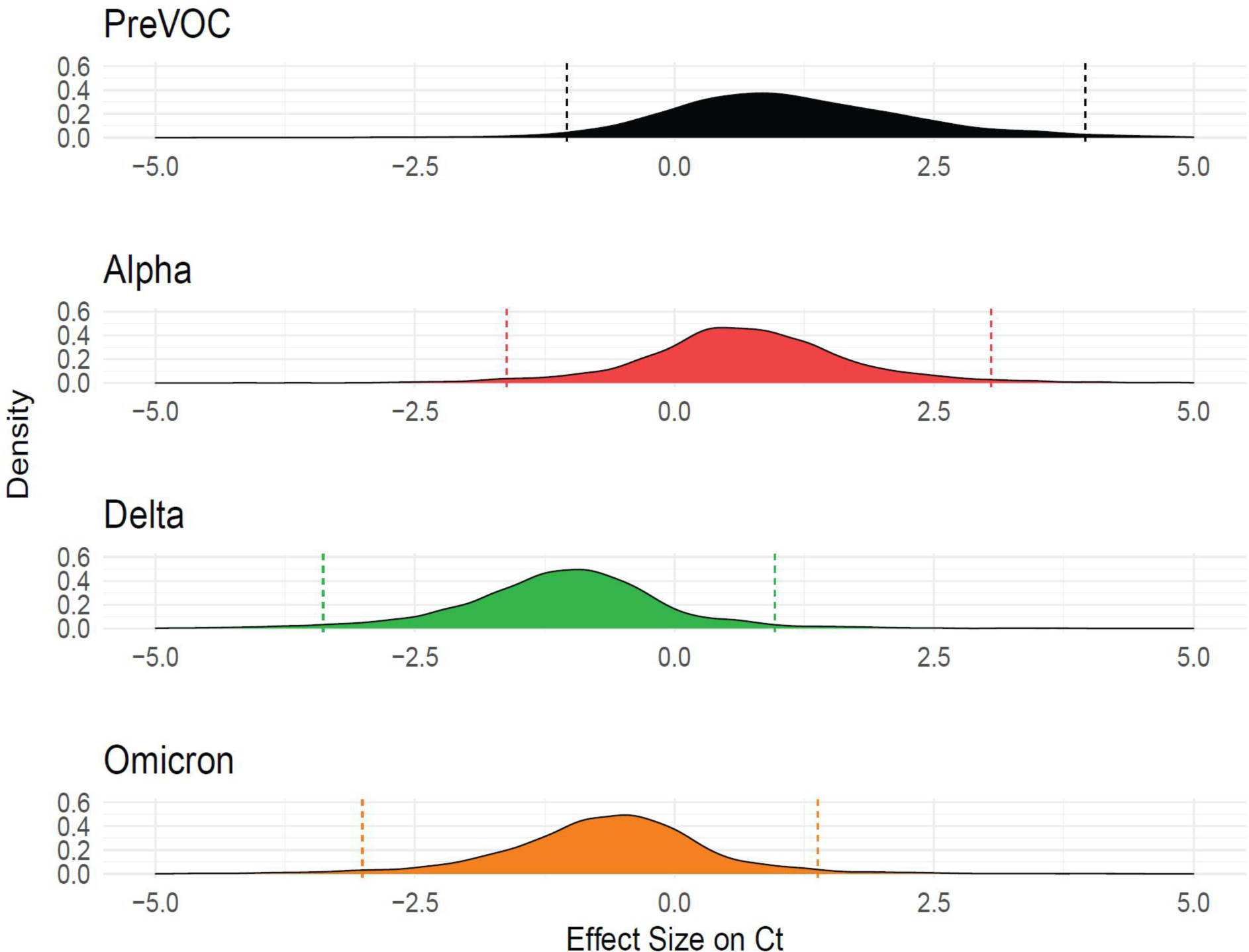
Delaware Valley SARS-CoV-2 surveillance data. (A) Effect size of variants on clinical qPCR assay Ct (compared to pre-VOC), adjusted for time during wave, specimen type, and qPCR machine as captured using a Bayesian regression model. The X-axis shows the marginal effect on Ct for each variant compared to preVOC. Note that a 1 cycle change in Ct is roughly equivalent to a 2-fold increase in RNA abundance under ideal PCR conditions. The Y-axis shows the posterior probability density generated from the Bayesian regression model.

We found that Delta specimens had Ct values that were estimated to be on average 1.77 Ct lower than Alpha (95% Credible Interval (CrI): 0.70, 2.89) and 0.43 Ct lower than Omicron (95% CrI: 0.15, 0.72). Omicron was estimated to have a Ct 1.35 cycles lower than Alpha (95% CrI: 0.28, 2.42). Delta was consistently found to have a lower Ct and thus higher numbers of RNA copies in infected human participants, followed by Omicron and Alpha. This finding is consistent with published studies of patient samples (43-46) and matches our findings that Delta and Omicron VOCs are most efficient at replicating in primary nasal cultures (**Fig 1**).

## Discussion

The COVID-19 pandemic has been marked with emergence of SARS-CoV-2 variants including Alpha, Beta, Delta and Omicron, which were designated as variants of concern (VOC) due to their heightened risk to the population. Infection with different VOCs have shown notable differences in patient health outcomes (47) but understanding of the differences in viral biology accounting for these differences is incomplete. A comparative mechanistic analysis of SARS-CoV-2 variants is necessary to delineate the virus-host biology driving pathogenesis and to help predict pathogenicity of future variants. Our findings parallel and provide mechanistic insight into retrospective human studies (6, 47) which showed that infections with Delta are most pathogenic, while Omicron is less pathogenic and instead selected for better replication (6).

Our findings focus attention on the differences in infections among SARS-CoV-2 viruses in primary nasal cultures and cell lines that facilitate technical aspects of some assays that are not feasible in primary nasal cultures. The cell lines used include A549 and Calu3, which allow us to compare infection in human lower respiratory tract derived cells. However, a caveat of such comparison is the effects of primary cultures versus cell lines derived from carcinomas to the observations. A strength of our study lies in analysis of infection in patient-derived primary nasal cultures which are the primary site of infection and model the upper respiratory tract (38). Unlike cell lines or animal models, our findings with the primary nasal cell model parallel human epidemiological studies in the comparison of infections by the different variants, in that, in both models Delta and Omicron were selected for efficient replication compared to other VOCs.

A consistent observation throughout our studies is that Delta infections are most damaging in both upper and lower respiratory model systems. This could be attributed to the polybasic furin cleavage site on the Delta spike protein, most basic among the VOCs, which can recruit proteases to facilitate spike cleavage and subsequent spread of infection. Despite sharing an identical furin cleavage site, Alpha and Omicron infections generate different levels of cleaved spike, cell to cell spread of virus, and cellular damage. Thus, our results indicate that factors in addition to the cleavage site play a role in these processes. Sequence and structure-based bioinformatics studies have proposed that mutations in Omicron encoding substitutions at the S1/S2 site may help evade recognition by proteases and prevent syncytia formation (48), which may explain our observation of less than expected levels of spike cleavage and propensity towards endosomal entry. However, more recent studies using nasal organoids have reported increased syncytia formation with the Omicron BA.5 subvariant compared to the first Omicron BA.1 variant used in our study (49, 50).

Differences in innate immune responses to infections by different VOCs have been studied extensively. We have shown that Omicron infection induces greater *IFNB* and *IFNL* gene response in primary nasal cultures, compared to the ancestral strain. However, in Calu3 cells, Beta infection elicited a stronger IFN response that surpassed that of infection with other VOCs. These variations may be explained by several factors. First, different cell types have different capacities for generating IFN responses. For example, it has been shown that nasal cultures have elevated basal levels of interferon gene expression and thus induction may appear to be blunted (38). Second, the level of innate responses may correlate with viral load in cells, and different VOCs replicate to different relative titers in different cell types.

Clinical studies of COVID-19 patient outcomes have proposed that patients with an early and robust INF response were more likely to have mild to moderate disease while patients with a delayed or blunted response were more likely to succumb to severe COVID (51, 52). The high induction of INF and ISGs with Omicron infections in our model would hypothesize that this induction may contribute to the milder disease associated with Omicron (5, 6). Recent studies also observed greater ISG induction with Omicron (53) and reduced IFN antagonism by Omicron as contributing to the milder disease (40). However, there are many confounding factors that warrant further experimentation.

Primary human nasal epithelia cultures grown at an air-liquid-interface are a powerful model to recapitulate coronavirus infections of the upper respiratory system (38, 54). Previous studies have validated SARS-CoV-2 infections in such models (16, 55, 56). Here we were able to characterize specific forms of cellular damage in ciliated cells by different VOCs. Immunofluorescence analysis shows that Delta is the most cytopathic VOC, followed by Alpha and Omicron. This order was also observed with levels of spike cleavage, suggesting that in nasal cultures Delta may spread by cell-to-cell fusion more than other VOCs. Analysis of TEER also identified Delta as most damaging to cell barrier integrity, followed by Omicron. However, further analysis suggests that Delta and Alpha caused the greatest loss of ciliary beating function. Therefore, while Delta infection compromises both functions, Omicron is detrimental to cell-barrier integrity more than ciliary beating, and Alpha infection diminishes ciliary beating but not cell-barrier integrity. This contrast highlights how the variants differentially affect mechanisms crucial for nasal epithelial functions at the cellular level that may correlate with variant-specific patient symptoms.

In primary nasal cultures, Delta and Omicron replicated to higher titers faster, while WA1 produced the lowest titer. In contrast, in the Calu3 cell line, replication of the earlier VOCs such as Alpha and Beta reached higher titers earlier. These results suggest that SARS-CoV-2 is evolving for efficient replication in the upper respiratory system. This observation concurs with clinical studies that report Omicron infections induce more upper-respiratory symptoms with reduced pathogenic manifestation (57, 58). These observations suggest future variants may continue to be selected for upper-respiratory infections and a general decrease in pathogenicity.

While the use of primary nasal epithelia model provides many advantages to our study there are some caveats to be aware of. The heterogeneity of cell types in the nasal epithelium compromise pathway and mechanistic analysis. While cell lines are used to circumvent these technical problems, this introduces confounding factors that are driven by cell line specific biology. The primary nasal cultures also lack an intact immune system that would be present in the animal.

The currently circulating SARS-CoV-2 virus is likely to continue to evolve. The findings in this study provide a comprehensive comparison of VOCs to date in cell culture, clarifying important differences in virus-host biology among SARS-CoV-2 variants affecting pathogenesis. These findings can be applied to understand and predict replication, spread and immunogenicity of future variants.

## Materials and Methods

### Viruses, replication curves and plaque assays

The following viruses were obtained from BEI resources, WA1/USA-WA1/2020 strain NR-52281 and Alpha NR-54000. Delta and Omicron were isolated from patient samples. All viruses were grown in VeroE6^TMPRSS2^ cells and titers were quantified by plaque assay on VeroE6 monolayer overlayed with 0.1% agarose and stained with 10% crystal violet (38). All viruses were sequence verified using the POLAR protocol using an Illumina NextSeq instrument with a 74 × 74 paired-end sequencing on a 150 cycle MID output cartridge (59). Infections were performed at MOI=0.1, unless otherwise specified, in serum-free DMEM for 1-hour, followed by refeed of cellular media for the duration of the experiment. For virus growth curve experiments, virus supernatant was collected at noted hours post infection (hpi) and quantified by plaque assay. For intracellular virus, cells were collected in DMEM and subject to 3 freeze-thaw cycles to release intracellular virus.

### Cell lines

VeroE6^TMPRSS2^ (African green monkey kidney) cells were maintained in DMEM (Gibco cat. No. 11965) with 10% fetal bovine serum (FBS), 100 U/ml penicillin, 100 ug/ml streptocymic, 50 ug/ml gentamicin, 1 mM sodium pyruvate, and 10 mM HEPES. Human A549^ACE2^ cells were cultured in PRMI 1640 (Gibco cat. No. 11875), 10% FBS, 100 U/ml of penicillin and 100 ug/ml streptomycin. Human Calu3 cells (clone HTB-55) were maintained in MEM media with 10% FBS.

### Genomes/PFU

Calu3 cells were infected with WA1 and VOCs at MOI 0.1, at 24hpi virus supernatant was collected. Infectious virus was quantified by plaque assay. Genomes were measured by extracting RNA from the supernatant and performing RT-qPCR with primers for SARS-CoV-2 nsp12 (RdRp) sequences.

### RNA extraction

Cells were lysed at indicated hours post infection in Buffer RLT (Qiagen cat. No. 79216) followed by extraction of RNA using the RNeasy Plus Mini Kit (Qiagen cat. No. 74004). Cell-free supernatant was lysed with AVL buffer (Qiagen cat. No 19073) and RNA was extracted with QIAmp Viral RNA Mini Kit (Qiagen 52904).

### RT-qPCR

The protocol for RT-qPCR was previously described and briefly outlined here (60). RNA was reverse transcribed into cDNA with a High Capacity cDNA Reverse Transcriptase Kit (Applied Biosystems). Target cDNA was amplified using specific primers, iQ SYBR Green Supermix (Bio-Rab) and QuantStudio 3 PCR system (Thermo Fisher). Host gene expression displayed as fold change over mock-infected samples was generated by first normalizing cycle threshold (CT) values to 18S rRNA to generate ΛCT values (ΛCT = CT gene of interest - CT 18S rRNA). Next, Λ (ΛCT) values were determined by subtracting the mock-infected DCT values from the virus-infected samples. Technical triplicates were averaged and means displayed using the equation 2^-Δ (ΔCT)^. Graphed values are the mean of biological triplicates of each condition and technical triplicates of each sample. Host gene expression was quantified with the following primers (forward sequence/ reverse sequence): IFNB (GTCAGAGTGGAAATCCTAAG/ CAGCATCTGCTGGTTGAAG), IFNL1 (CGCCTTGGAAGAGTCACTCA/ GAAGCCTCAGGTCCCAATTC), OAS2 (TTCTGCCTGCACCACTCTTCACGAC/ GCCAGTCTTCAGAGCTGTGCCTTTG), IFIT1 (TGGTGACCTGGGGCAACTTT/ AGGCCTTGGCCCGTTCATAA) and 18S rRNA (TTCGATGGTAGTCGCTGTGC/ CTGCTGCCTTCCTTGAATGTGGTA). Virus genomes were quantified in reference to a standard curve, with primers for SARS-CoV-2 genomic nsp12/RdRp (GGTAACTGGTATGATTTCG/ CTGGTCAAGGTTAATATAGG).

### RNA Sequencing

Nasal epithelial cells from 4 donors were pooled together into air-liquid interface cultures. Mock infections or infections with the indicated viruses were performed in triplicate at MOI=0.1. Cells were lysed at 96 hpi using RLT Plus buffer and total RNA was extract using Qiagen RNeasy Plus Mini kit (cat. No. 74004). Samples were sent to Azenta Life Sciences for RNA sequencing with Illumina HiSeq PE 2x150. Read quality was assessed using FastQC v0.11.2 (61). Raw sequencing reads from each sample were quality and adapter trimmed using BBDuk 38.73 (62, 63). The reads were then mapped to the human genome (hg38 with Ensembl v98 annotation) using Salmon v0.13.1 (64). Differential expression between mock and infected experimental conditions were analyzed using the raw gene counts files by DESeq2 v1.22.1 (65). Volcano plots were generated using EnhancedVolcano v1.14.0 (66), with highlighted interferon stimulated genes (ISGs) being selected from the Molecular Signatures Database HALLMARK_INTERFERON_ALPHA_RESPONSE gene list (67).

### Data Availability

Raw and processed RNA-seq data for all infection conditions will be deposited in the Gene Expression Omnibus database prior to publication.

### Graphical visualization, quantification, and statistics

Graphs were generated using Prism software. Statistics were also performed with Prism software with specific tests stated with each experiment. Plaque area quantification, syncytia area quantification and western blot quantification were performed with ImageJ software.

### Protease treatments

Nafamostat (20uM) was used for protease inhibition (68). Calu3 cells were pre-treated for 2 hours with appropriate concentrations of drug in the cell media. Infections were performed at MOI=0.1 for 1hour, after which inoculum was replaced with Calu3 media plus drug for 1 additional hour of post-treatment. After this cell media was replaced with drug-free media for the remainder of the experiment. For positive control, drug concentration was maintained in the media for 4 hours of post-treatment. For negative control, no drug was included in the media and DMSO media (dimethyl sulfoxide, Thermo Scientific cat no J66650) was used instead. Supernatants were collected at 16hpi for quantification by plaque assay. Percent inhibition was calculated as fraction of virus titer after drug treatment over virus titer without drug treatment.

### RNase L degradation assay

RNA integrity was analyzed on a chip with Aligent 2100 Bioanalyzer using the Aligent 196 RNA 6000 Nano Kit (Cat #: 5067-197 1511)

### Western Immunoblot

Cells were washed in PBS and harvested in lysis buffer (1% NP-40, 2mM EDTA, 10% glycerol, 150mM NaCl, 50mM Tris-HCl, protease inhibitor (Roche complete mini EDTA-free protease inhibitor), phosphatase inhibitor (Roche PhosStop easy pack). After 20 minutes of lysis on ice, samples were centrifuged to remove cell debris. Lysates were denatured at 95C for 5 minutes and stored for analysis. Protein lysates were separated on 5-15% SDS-PAGE gradient gel and transferred onto a PVDF membrane. Membrane blots were blocked with 5% nonfat milk or 5% BSA, and probed with appropriate primary antibodies overnight at 4C, and secondary antibodies for 2h at room temperature. Blots were exposed with chemiluminescent substrate (Thermo Scientific Cat. No. 34095 or 34080). Blots were stripped (Thermo Scientific cat no 21059) and reblotted as needed.

### Primary human nasal cultures

Sinonasal cells were obtained from patient donors with informed consent, per protocol approved by the University of Pennsylvania Institutional Review Board (protocol #800614). A detailed protocol was described previously (38, 69). Briefly, specimens were dissociated and grown to 80% confluence in in PneumaCult™-ALI Medium (STEMCELL Technologies 05001) supplemented with heparin (STEMCELLl Technologies 07980) and hydrocortisone. Once cells reach confluency in 0.4 µM pore transwell inserts, the apical growth media is removed and the basal differentiation media is replaced every 3-4 days for a minimum of 4 weeks. Prior to infection epithelial morphology and cilia beating is confirmed visually by microscopy.

### Transepithelial electrical resistance (TEER)

TEER measurements were obtained with the EVOM apparatus, in PBS supplemented with calcium and magnesium. TEER measurements were obtained for all nasal ALI culture transwells pre-infection (0hpi) and post-infection (48hpi and 96hpi). ΔTEER is the difference in TEER from 0hpi to 48hpi, and 48hpi to 96hpi.

### Cilia beat frequency and beating cilia

Live microscopy movies of nasal cultures were obtained with a 20x objective on a brightfield microscope. Movie segments were analyzed on SAVA system to obtain cilia beat frequency (70). Beating cilia was obtained with single point analysis. Graphed values are an average of three nasal cultures per condition and four regions of interest per culture.

### Immunofluorescence

A549^ACE2^ cells were seeded on coverslips and infected at confluency at MOI=0.1. At 24hpi samples were fixed with 4% paraformaldehyde for 30 minutes, followed by permeabilization with 0.5% trypsin for 10 minutes. Samples were blocked with 1% BSA, followed by incubation with primary antibodies for 2 hours at room temperature, and secondary antibodies for 1 hour at room temperature. Antibodies used include DAPI, Nucleocapsid (1:500, gift from Dr. Tony Schountz, Colorado State University, Fort Collins, CO, USA), Alexa-488 goat anti-mouse (1:1000, Thermo Scientific cat no A11011). Images were obtained with a 60x objective on a Nikon Ti-8 brightfield microscope. For nasal cultures the images were captured on a confocal microscope and displayed as overlay projections.

### Ct analysis of Delaware Valley surveillance

Samples were collected from the Delaware Valley under University of Pennsylvania IRB protocol no. 823392, CDC BAA 200-2021-10986 and CDC 75D30121C11102/000HCVL1-2021-55232. RNA samples were analyzed by whole genome sequencing as described previously (42, 71). Ct values were obtained from patient records. Clinical viral RNA level measured by cycle threshold (Ct) was predicted using a Bayesian regression model as implemented in BRMS using a thin plate spline for time since variant first detected and random effect of qPCR machine, specimen type, and variant of sample.

*Ct_i_* ~ *s*(*t_i_*) + *λ_i_* + *α_v_* + *β_m_*

Where *Ct_i_* is an estimate of a sample’s Ct value. *s*(*t_i_*) is a thin plate regression spline calculated to account for variations over time for all variant’s introduction. λ*_i_* is the effect on sample location on either upper or lower respiratory tract. *α_v_* is the effect for a given variant, and *β_m_* is the effect for a given qPCR protocol.

2,722 SARS-CoV-2 positive participants had clinical Ct data collected and were sequenced through whole-genome sequencing from 2/15/2021 to 7/18/2022. The supplemental table contains metadata and sequence accessions used in this analysis. One of six qPCR protocols were used (Cepheid GeneXpert, Cobas8800, Cobas Liat, DiaSorin MDX, Saliva COVID, or ThermoFisher Amplitude). One of five variant categories were assigned from whole genome sequencing (Alpha, Delta, Omicron, other variant, pre-variant of concern).

## Acknowledgements

We thank Dr. Paul Planet for the Delta variant sample, Peter Hewins and Dr. Kellie Jurado for the Beta variant, and Dr. Pei Young Shi for the icWT and icPRRA deletion viruses. We also thank Drs. David W. Kennedy, James N. Palmer, Nithin D. Adappa, and Michael A. Kohanski for aid in the collection of nasal tissue for establishing primary nasal epithelial cultures. This work was supported by National Institutes of Health grants R01 AI140442 (SRW) and R01AI169537 (SRW&NAC); Department of Veterans Affairs Merit Review 1-I01-BX005432-01 (NAC&SRW); Centers for Disease Control and Prevention contract award BAA 200-2021-10986 and 75D30121C11102/000HCVL1-2021-55232 (FDB). NST was supported in part by T32NS43126-18. This work was also supported by philanthropic donations to the Penn Center for Research on Coronaviruses and Other Emerging Pathogens (SRW&FDB) and the Penn Center for AIDS Research (P30-AI045008).

---

**Supplemental Figure 1:**
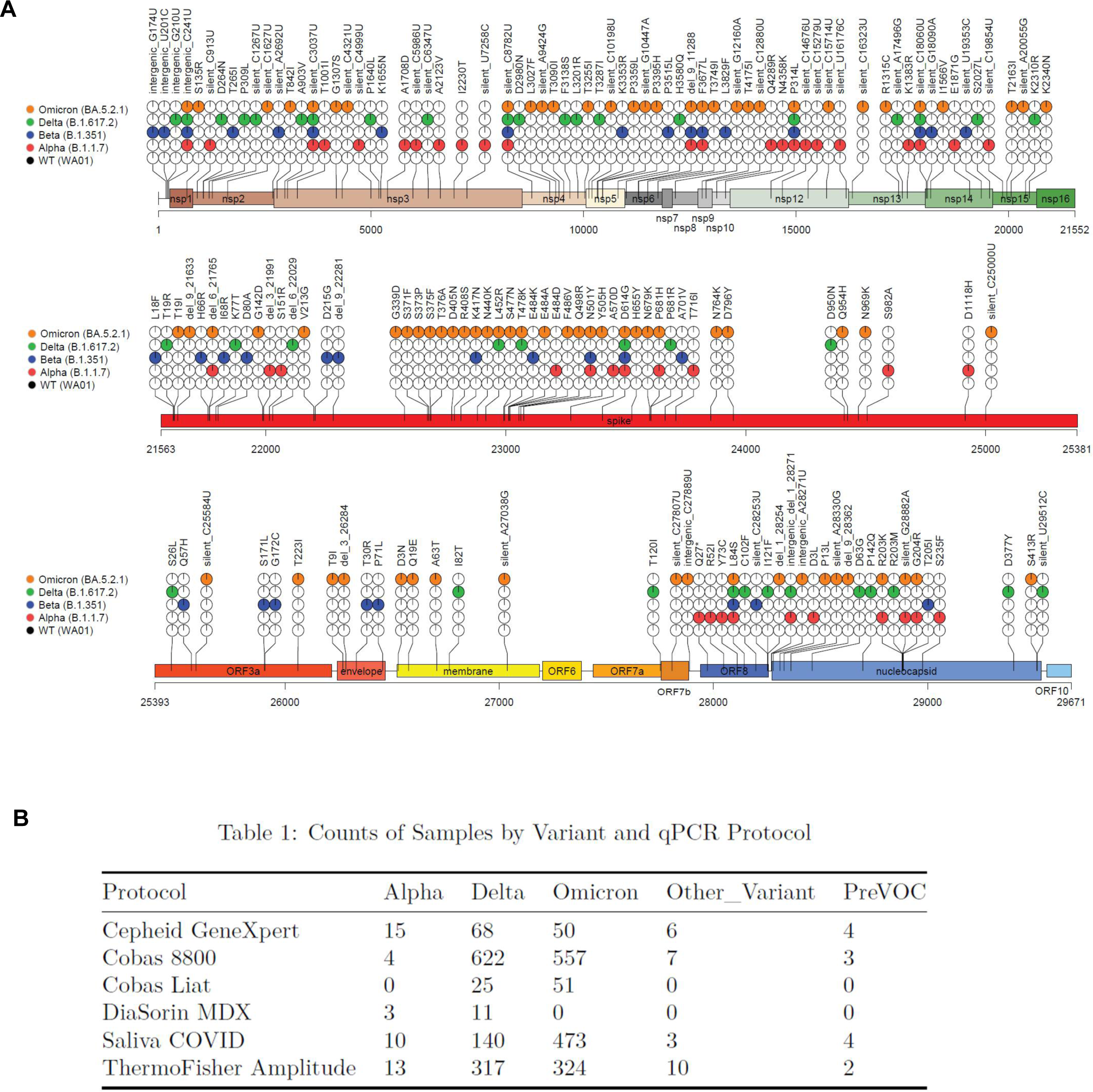
SARS-CoV-2 infections in cell lines. (A) SARS-CoV-2 stocks were harvested from VeroE6^TMPRSS2^ cells after 2 passages and RNA was collected from infected cells and sequenced. Viral sequences were aligned against the WA1 genome sequence for reference. Amino acid substitutions are indicated with colored circles, black for WA1, red for Alpha, blue for Beta, green for Delta and orange for Omicron. The fraction of sequences with indicated mutation is denoted as pie slice within the circle. (B) Different analytical technologies and PCR protocols used to measure the graphed Ct values throughout the endemic.

**Supplemental Figure 2:**
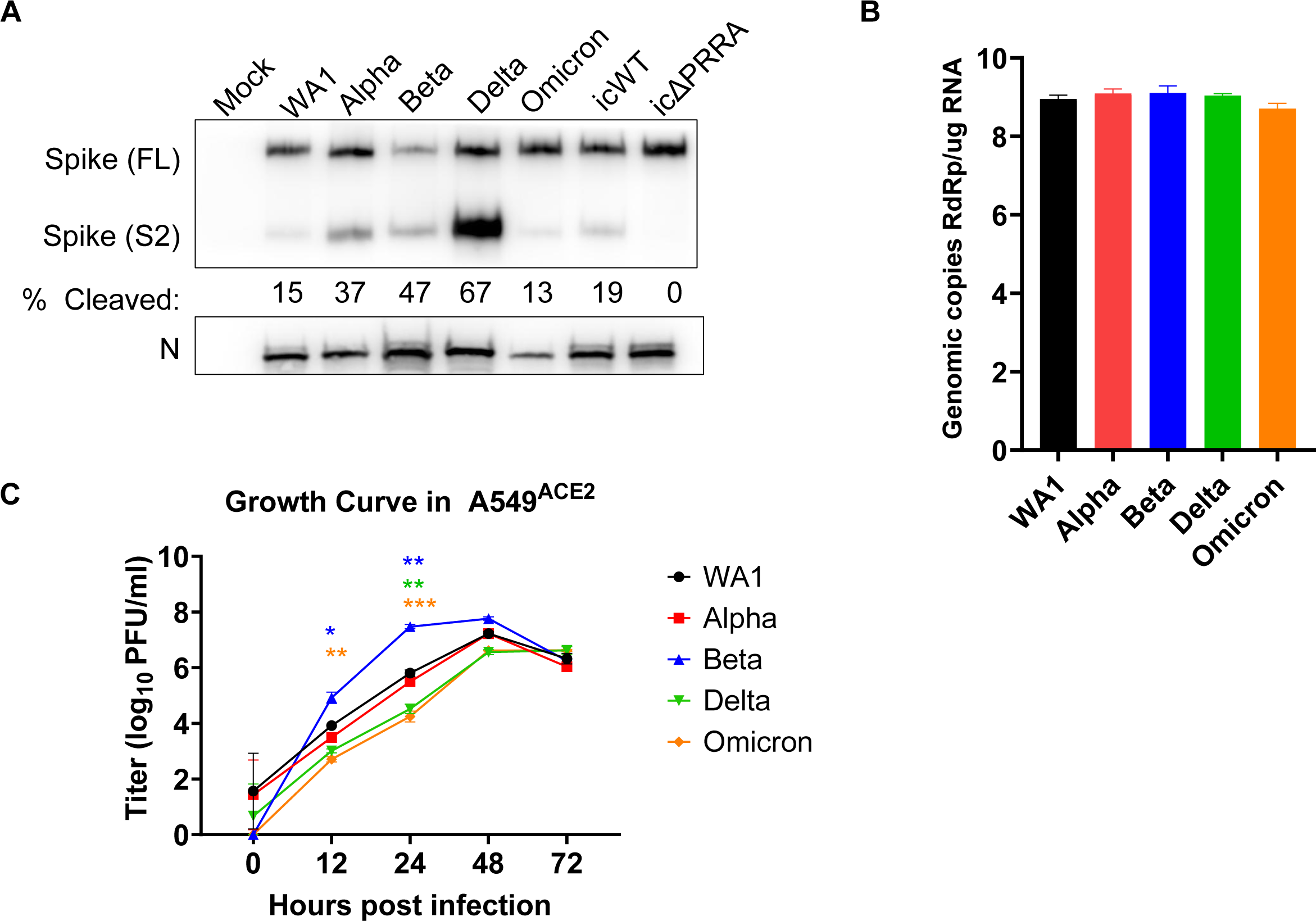
SARS-CoV-2 infections in lower respiratory cell lines. (A) VeroE6^TMPRSS2^ cells were infected at MOI=1 and lysed for protein at 48 hpi. Proteins were analyzed by western blot and probed with antibodies against nucleocapsid (N) and spike (S2). Band intensities from 3 independent western blots were quantified to measure the fraction of cleaved spike protein. The percent of cleaved spike was calculated as the fraction of cleaved spike (S2) over the sum of full length (FL) and cleaved (S2) spike. (B) Calu3 cells were infected with SARS-CoV-2 viruses (MOI=0.1) and RNA was collected at 24 hpi. RT-qPCR was performed with primers specific for SARS-CoV-2 nsp12 RdRp sequence to quantify virus genomes. (C) A549^ACE2^ cells were infected (MOI=0.1) and supernatant was collected at indicated timepoints to quantify infectious virus by plaque assay. Graphed values represent mean with standard deviation, and statistics were performed with ordinary two-way ANOVA with multiple comparisons for VOCs versus WA1 within a time point, adjusted P-values: *P<0.05, **P<0.005, ***P<0.0005.

**Supplemental Figure 3:**
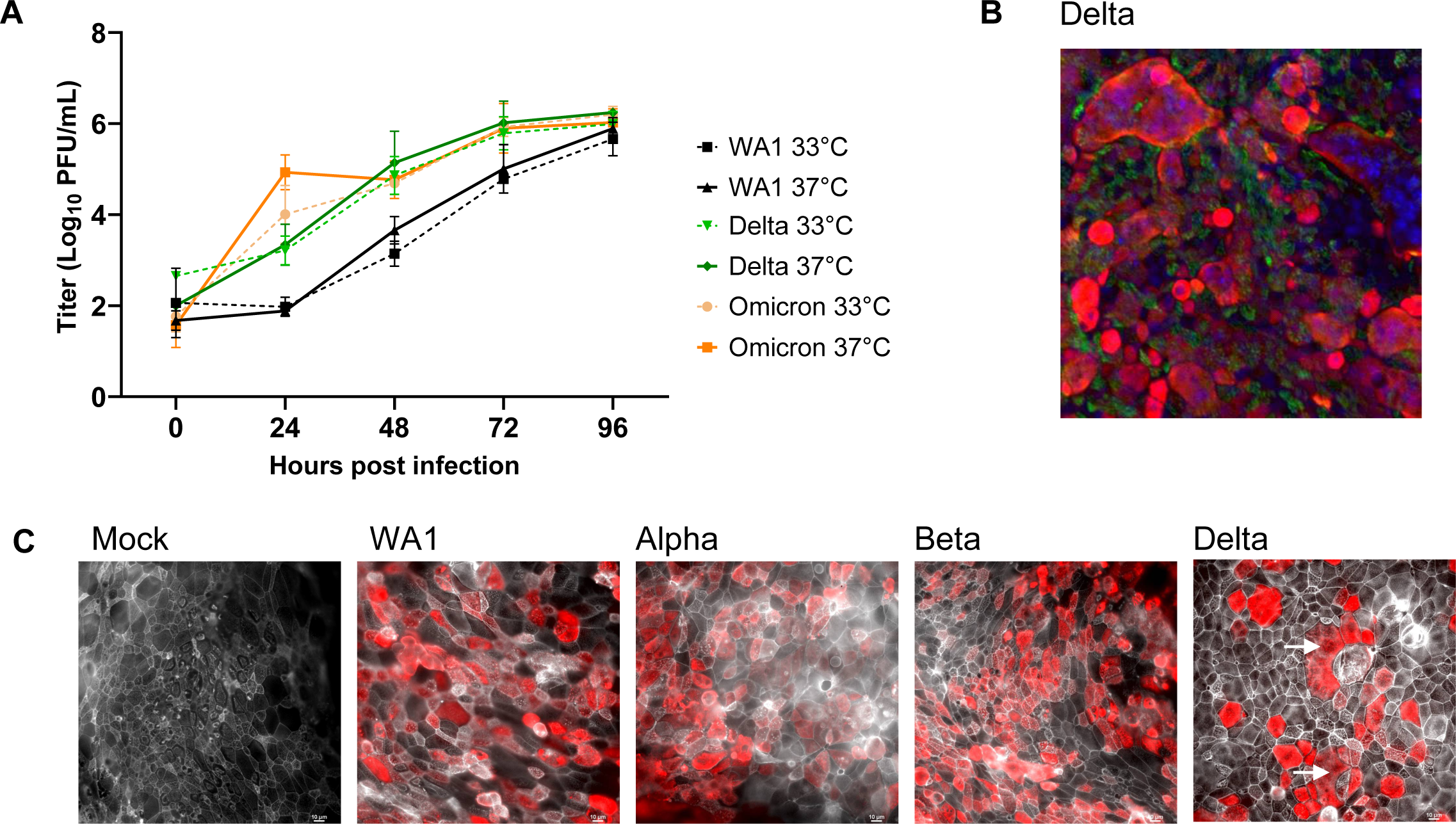
WA1 and VOC infections in nasal cultures. (A) Nasal cultures (pooled from 4 donor cells) were infected (MOI =0.1) and incubated at 33°C or 37°C. Apically shed virus was collected at the indicated time points and infectious virus was quantified by plaque assay. Growth curves at 33°C are indicated with dashed lines, and 37°C with solid lines. Graphed values represent mean with standard deviation and statistics were performed with ordinary two-way ANOVA with multiple comparisons for VOCs versus WA1 at the two temperatures within a time point. Values did not reach statistical significance. (B) Nasal cultures were infected at MOI=0.1; at 96 hpi samples were fixed and processed by immunofluorescence for DAPI (blue), β-tubulin (green) and SARS-CoV-2 nucleocapsid (red). (C) Infected and mock cultures (also depicted in Fig 7D) were fixed for immunofluorescence and confocal imaging with antibodies to label N (red) and phalloidin (white). Arrow indicates syncytia-like clusters. Images are representative of infections in multiple donor nasal cultures.

